# Effects of joint invasion: how co-invaders affect each other’s success in model food webs?

**DOI:** 10.1101/2024.01.02.573872

**Authors:** Ágnes Móréh, Ferenc Jordán, István Scheuring

## Abstract

While there is considerable research on interactions between invasive and native species, as well as on the impact of invasive species on the resident community, less focus has been placed on exploring the relationship and interactions among invasive species themselves. Nevertheless, it is widely acknowledged that invasive species can have either positive or negative effects on one another’s success, in addition to neutral outcomes. In the present theoretical study, we compared the success of two invasive non-native species in two scenarios: when they invaded the resident food web separately and simultaneously. We investigated the correlations between their direct and indirect ecological relationships and the topological positions of them in the food web, with the varying outcomes of joint invasion. Using the allometric bioenergetic model (ABM) for dynamical simulations, we detected the success of invasion (presence or absence of invaders) and the direction of their biomass change comparing separated and simultaneous invasion scenarios. We studied the relationships among these variables after detailed numerical simulations with variable key parameters of the model.

We found that direct and indirect ecological relationships between the two invaders are significantly modifies the invasion scenarios: the predator-prey relationship increases the probability of invasion success for both invaders, but the equilibrium biomass of at least one of them is more likely to be reduced than in separate invasions. The trophic cascade or competitive relationship between them during simultaneous invasion also affects their success rate, with the former having a positive effect and the latter a negative one. Further, we found that higher trophic level and lower betweenness centralities of the invaders reduces the likelihood of invasion success regardless of the presence or absence of another invasive species. The results of the study can be tested experimentally in micro- and mesocosms.

**HIGHLIGHTS:**

- In a joint invasion, invaders can influence each other’s success
- Predator-prey relationships between invaders increase the joint invasion success
- Competition increases the failure of at least one invader
- The trophic cascade between invaders increases the joint invasion success
- Higher trophic levels or lower betweenness centralities of the invaders increase the probability of unsuccessful joint invasion

**GRAPHICAL ABSTRACT:** 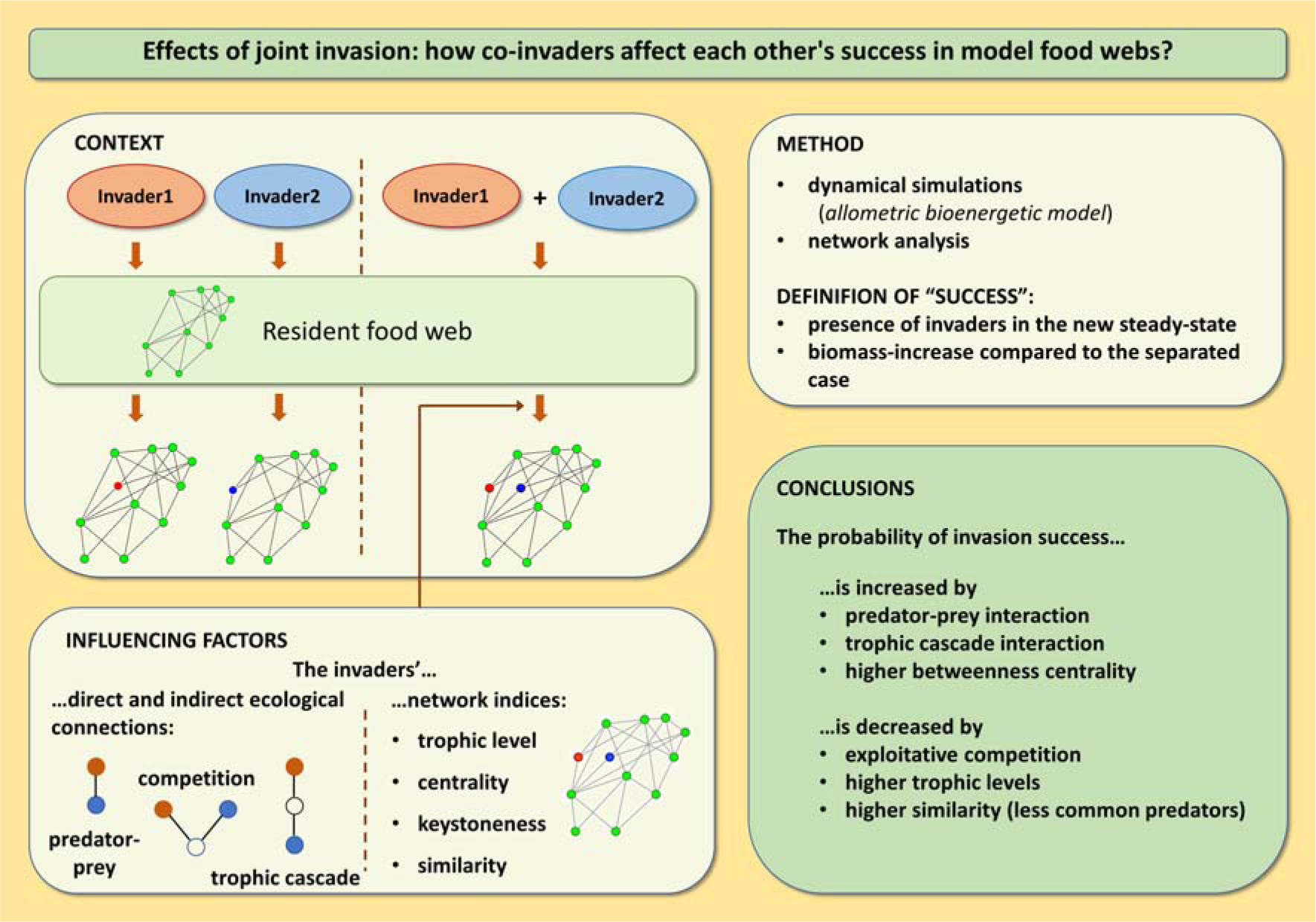

## 1. INTRODUCTION

Studying biological invasions is crucial as it helps us understand the far-reaching ecological and economic consequences of non-native species establishing themselves in new habitats. Although the majority of research tends to concentrate on the single invasive species, there is an increasing significance in grasping the interactions between the invaders that share habitats they have colonized together (Kuebbing et al., 2013). Despite of the widespread co-occurrence of multiple invaders sequentially or in parallel, little is known about their influence on each other, or about their combined ecological impacts on native communities (Dénes et al., 2018).

Invasive species establish difficult-to-predict relationships with both indigenous and non-indigenous species, potentially reshaping the configuration and progression of communities (Hewitt and Huxel, 2002). By means of consumption or competition, they might lead to decreases in populations if affected resident species are unable to adapt their defensive tactics or resource utilization (Dueñas et al., 2021). Nonetheless, the impact of invasive species frequently hinges on the characteristics and ecological roles of the impacted species in relation to the invasive species (David et al., 2017).

Even though investigating biological invasions constitutes a vast field, the literature on bi- or multi-species invasions is notably limited. This limitation is particularly pronounced in the realm of theoretical modeling. One of the exception is the study by Hewitt and Huxel (2002) where the focus is on how the number or initial density of one or more invasive species influences both their success and the resilience of the native ecosystem, however, they didn’t discuss the reciprocal impacts of two invasive species on one another or on the whole ecological network. In reality, invasive species engage in interactions with both native species and with other invaders within ecological networks, leading to a diverse range of direct and indirect effects within the community (Simberloff et al., 2013). While theoretical studies are practically missing in this area the joint effects of two (or more) invasive species on each other or on the ecosystem are investigated mainly through mesocosm or field experiments (Cope and Winterbourn, 2004; Johnson et al., 2009; Preston et al., 2012; Braga et al., 2020). The potential robust trends are explored as well by using meta-analyses of these studies (Kueffer et al., 2013; Jackson, 2015). They found that i) invasive animals generally tend to have neutral or negative effects on each another, and ii) their cumulative effects on the biodiversity, e.g. on total community abundance or on survival of the native system, are generally less pronounced than what would be expected in additive scenarios, while synergistic effects are rare. However, they note that some case studies, field or mesocosm experiments often give contradictory outcomes (e.g. Crain et al., 2008), and there is no getting around the fact that invasive species can have positive effects of on each other, as highlighted by the Invasional Meltdown Hypothesis (Simberloff and Holle, 1999).

Available evidence suggests that when more invasive species appear together, complex and unpredictable interactions among the species can influence invasion outcomes (Russell et al., 2014). In this paper, our aim is to compare the effects of two invaders in two different settings: when they invade together and separately into a resident community. Actually, the present study focuses on a pivotal aspect: the mutual influence of the invaders on their relative invasion success. The success of an invasive species is determined by its ability to establish and maintain a viable population within the resident community (Kolar and Lodge, 2001; Blackburn et al., 2011). It is plausible, for example, that both invaders succeed when they enter the native system individually, however, in the presence of each other, one may not survive or both could become extinct. Another possible scenario is that invasion is not possible alone, but this invader succeeds in association with another invader. Interactions can either facilitate or hinder the invaders’ spread within the community.

In addition to analyzing the frequency of different outcomes of invasion and comparing their impacts in separate and joint scenarios, our primary focus was to identify the most influential factors that may affect the invasion outcome. It is highly probable and testable that the position of the invading species within the food web influences their impact on each other. It may matter, for example, what trophic levels they invade at, how different they are in terms of their centrality in the food web, how different their prey or predators are, or simply what are the nature of the ecological relationship between them (predator-prey, trophic cascade or competition). These ecological connections can vary based on the identities of the invaders and their positions within the food web, thereby leading to cumulative effects of multi-species invasions within the broader context of the food web (Jackson et al., 2017; Zhang et al., 2019).

In summary, we explore the following questions:

`o` How do the invaders interact? When invading species enter an established food web together, how often do they help, hinder or have no effect on each other?
`o` Do simultaneous invasion outcomes depend on direct and indirect ecological interactions between invaders?
`o` How do different topological indices, like trophic level, centralities or keystoneness, have an impact on the invasion outcome of invaders within the food web?
`o` If only one invader is successful, which one is it? Furthermore, which positions or topological indices could indicate its’ success?

## 2. METHODS

### 2.1 Generating „native” food webs

As a first step, we created a collection of stable food webs with sizes between 10 and 20 nodes (*S*). By generating 10 networks for each node size, we obtained a total of 110 food webs. During this process, we defined some topological requirements that all networks had to fulfil. These constraints were determined by using literature data (Dunne et al., 2002; Hall and Raffaelli, 1991; Williams and Martinez, 2004) and by topological analysis of real (aquatic) networks already studied (Colléter et al., 2015; Heymans et al., 2014). The data of this analysis are summarized in the Table S1 in the Supplement 1. Previous studies have shown that the connectivity of the food web influences invasion success (Romanuk et al., 2009; Baiser et al., 2010). Therefore, to exclude this effect, we set connectivity (*C*) to a fixed value of 0.2, which is a reasonable value for most studied food webs for this size range (Heymans et al., 2014).

#### Construction of the native food webs

Beside applying the fixed connectivity rule we applied the following topological rules:

- The number of basal and top species varies between 1 and 4 selected from an even distribution.
- The maximum of the trophic levels is 5.
- Loops are excluded, all consumers can only prey on lower trophic levels.
- Omnivory is not excluded, instead, based on our topological anaysis of the real food webs (Table S1.), we define a maximum range for consumers from which we draw their prey and predators. Actually, for an *i*th consumer, this is *i* ± 0.75*S* (*S* refers to the node number), corresponding to preys and predators, respectively. For basal species, this range was slightly narrower, with predators selected from the range *i*_basal_ + 0.5*S*.
- Because of the fixed connectance, the number of nodes determines the number of links (*L* = *S*^2^*C*). The maximum number of links is defined by the average number of links per species 2*L*/*S* (Dunne et al., 2002).

#### Testing the constructed food web

If a food web met the structural requirements, we tested its stability using dynamic simulations (see details below). A network was considered suitable for invasion analysis if

- the system performed stable fixed point in steady state, and
- all species were coexisting (none of the resident species has gone extinct).

### 2.2 Food web dynamics

In the process of network generation and during the simulation of invasion, we used a modelling framework based on an allometric approach. The so-called *allometric bioenergetic model* (ABM) is a well-defined and widely used modelling approach in studies related to various types of food webs. As we applied the model and its parameters without any modifications, this summary represents an excerpt derived from the model description found in (Brose et al., 2017).

In the allometric coefficients-updated consumer-source model (Brown et al., 2004), a set of differential equations describes the changes in relative biomass density (*B_i_*) of basal and consumer species:

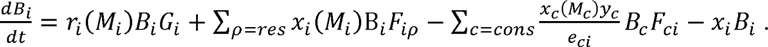

The first term on the right-hand side is applicable only to the basal species and does not apply to the consumers. Here, *r_i_* represents the mass-specific maximum growth rate of species *i*, and *G_i_* is the term for the logistic net growth rate (*G_i_* = 1 - (*B_i_* / *K_i_*), where *K_i_* denotes the carrying capacity associated to species *i*). Additionally, *x_i_* represents the mass-specific metabolic rate, and *y_i_* denotes the maximum consumption rate relative to its metabolic rate. Summations are for all resources (res) and consumers (cons). Finally, *e_ij_* represents species *i* assimilation efficiency when consuming species *j*.

The nonlinear functional response terms describe the consumption rate that is actually realized when species *i* consumes species *j*:

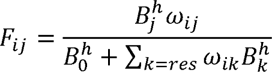

where *ω_ij_* represents the relative consumption rate of species *i* when consuming species *j*, *B_0_* is the half saturation density, and *h* is the Hill-coefficient. For consumers with *n* resources, we use uniform relative consumption rates (*ω_i_* = 1/*n_i_*). We set *h* = 1.5, which compromise the type II response (*h* = 1) and type III functional response (*h* = 2).

The core idea of the model lies in the negative quarter power law relationships governing the biological rates of *production* (*R*), *metabolism* (*X*), and *maximum consumption* (*Y*) in relation to the body mass of the species:

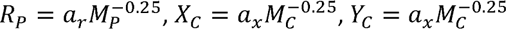

where *a_r_*, *a_x_* and *a_y_* are allometric constants, *C* and *P* indicate consumers’ and primary producers’ parameters, respectively (Brown et al., 2004; Yodzis and Innes, 1992). By setting the growth rate of the basal species to unity (*r_basals_* = 1), we establish the time scale for the system. The metabolic rates of the species are then normalized relative to this time scale:

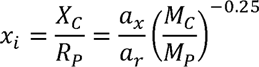

while the maximum consumption rates are normalized by the metabolic rates:

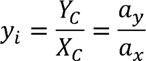

On the other hand, we take advantage of the observation that in (especially marine) food webs, predators usually exhibit larger body sizes compared to their prey. The predator-to-prey biomass ratio serves as a crucial metric for understanding trophic structure and community dynamics (Trebilco et al., 2013; Perkins et al., 2022). We introduce a constant predator-prey body mass ratio (Z), where Z = 10^2^ means that the predator is 100 times heavier than its prey (Brose et al., 2006). Consequently, we establish a relationship between the body mass and the trophic level (TL) of a species, represented by the expression: *M_C_* = *Z^TL^*.

In this context, the body masses of all predators are expressed relative to the body mass of the basal species, ensuring that the results are independent of the specific basal species chosen. This approach allows us to account for the variation in body mass with increasing trophic levels, providing insights into the size structure and dynamics of the predator-prey interactions within the food web.

In this deterministic model the network structure defines the dynamics of the populations, as the trophic levels of the species define their body mass, which defines the mass-dependent parameters, as shown in Fig 1. In the course of the simulations, the mass-independent parameters are kept constant and fixed based on previous work (Brown et al., 2004). The parameters used and their fixed values are listed in Table 1.

**Fig. 1.**
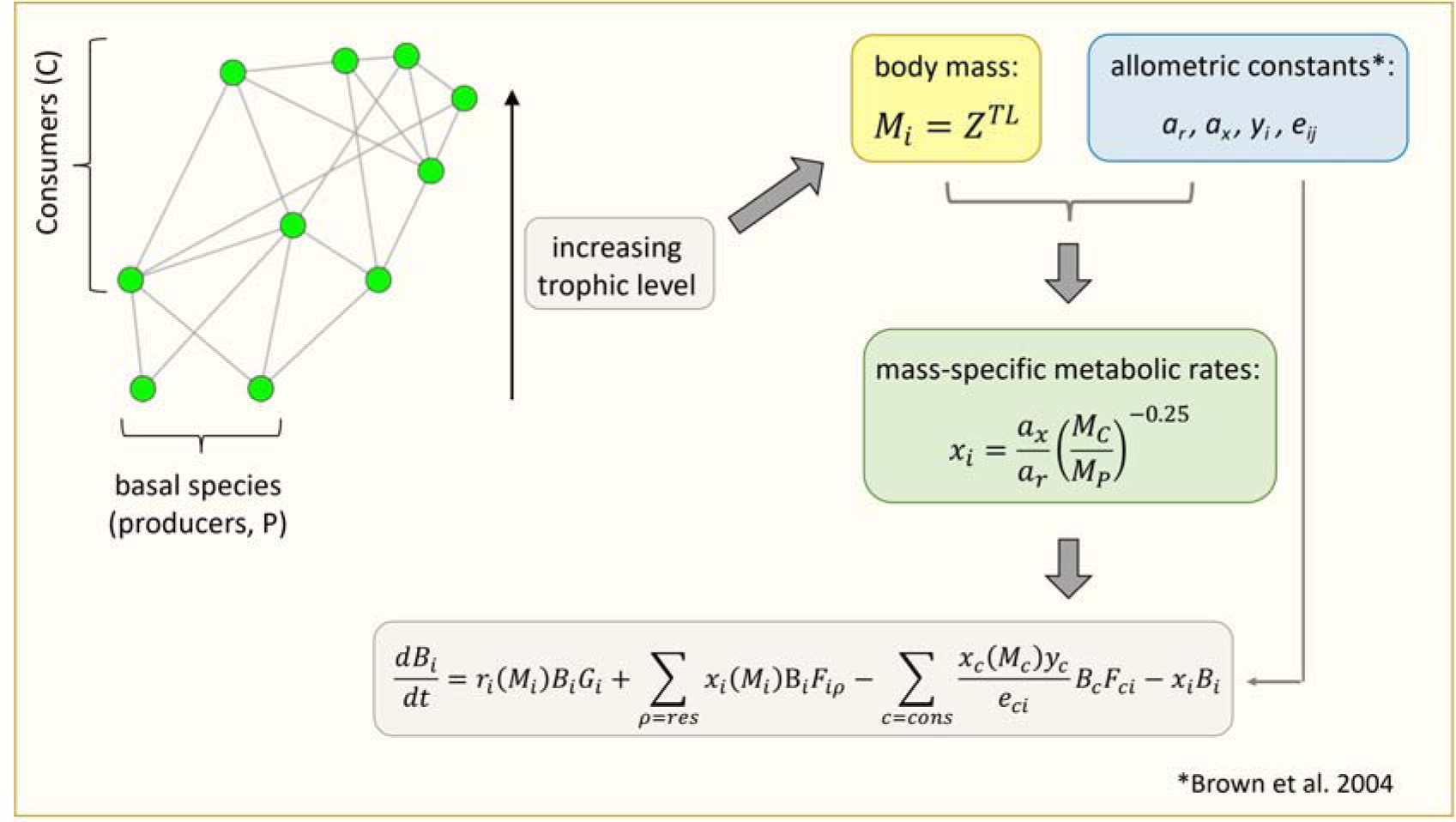
Outline of the process for creating the food web model. The food web topology, the metabolic and allometric parameters determine the dynamics (for more details see in the main text).

**Table 1.** Model parameters; C: consumers; P: primary producers; n_i_: number of i’s resources; a_x_, y_i_ and e_ci_ represent the mean of the constants given for vertebrate/invertebrate and herbivorous/carnivorous species

| **Parameter** | **Description** | **Fixed value** |
| --- | --- | --- |
| $B_i$ | biomass of $i$ | $B_{init} = 1$ |
| $r_i$ | mass-specific growth rate | 1 |
| $K_i$ | carrying capacity | 1 |
| $Z$ | consumer-resource body mass ratio (BMR) | 100 |
| $x_i$ | mass-specific metabolic rate | $\frac{a_x}{a_r} \left( \frac{M_C}{M_P} \right)^{-0.25}$ |
| $a_r$ | allometric constant | 1 |
| $a_x$ | allometric constant | 0.6 |
| $y_i$ | max. consumption rate | 6 |
| $e_{ci}$ | assimilation efficiency of $c$ when consuming $i$ | 0.65 |
| $B_0$ | half-saturation biomass | 0.5 |
| $\omega$ | relative consumption rate | $1/n_i$ |
| $h$ | Hill-coefficient | 1.5 |

### 2.3 Simulation of invasion

To simulate invasion, we expanded the food webs we created previously by adding the invading species to the web depending on the specific scenario being examined (separate or joint invasion). To assign the links for the invaders, we randomly selected them based on the same criteria used during the generation of the resident food web described above. Moreover, considering the characteristics of the model (i.e., the logistic growth term of the basal species ensures their constant presence, so basal invaders always invade successfully), we specified that only consuming species could act as invaders and that they must have at least one prey.

Our general question is how the outcome of the joint invasion modified, compared to when they appear separately in the system. However, for practical reasons, we reversed the process of expanding the generated food webs: first, the simulation for the joint invasion is proceeded, and subsequently, in the next round, we ran the simulation again with the same invaders but with the previously drawn links reintroduced separately back into the resident networks. This reverse ordering was crucial to investigate any potential predator-prey relationships that might arise between the two invaders during the analysis. In this manner, such relationships appeared randomly during the link draw in the joint invasion simulation, rather than requiring artificial creation of them afterward. This approach allowed us to effectively examine the distinct impacts of the invaders while considering the potential interactions between them in a more natural manner.

Although the invaders appeared almost simultaneously in the simulation with a very short time lag, for practical reasons, they were labelled as *I1* and *I2* denoting the order of appearance. The pairing was done by inserting *I1* right above the basal species (*I1*_Nbasal+1_), fixing its position, and then inserting *I2* next to it in sequence for all the levels above I1 (*I2*_I1+1…S+2_). For instance, if a resident network contained *S* number of species with three basals, *I1* would be the first invader as species #4 and would be systematically paired with I2 (#5 … #S+2). After each “round”, *I1* moved up one position and was paired again with the *I2* “above” it until both invaders were at positions #S+1 and #S+2. This technical solution ensures that invader-pairs emerge systematically across the entire consumer spectrum within a network.

For each pair, 10 different sets of links were randomly drawn for each invader. If the drawn combination resulted in a stable state (stable fixed point of the dynamics) at the end of the simulation, the combination was kept, otherwise it was discarded and a new one was drawn. More complex stable state (limit cycle, chaos) was not observed. This process yielded a grand total of 121000 invasion scenarios that could be tested, involving 110 resident food webs.

The generation of the resident food webs and the simulation of the invasions were performed with a C-code using the *CVODE* differential equation solver (Hindmarsh et al., 2005). We used *R* for evaluating the results as well as conducting the statistical analyses (R Core Team, 2020). The statistical tests used are summarised in the Supplement 1.

### 2.4 Influencing factors

In order to quantify the position of invaders in the resident food web, we calculated several network indices expressing structural and positional importance:

- *Trophic level* (*TL*) - we used the prey-averaged trophic level metric (Williams and Martinez, 2004) which is calculated as 1 plus the average trophic level of all the food sources for a given consumer;

#### - Centrality indices (Wasserman and Faust, 1994)

`o` *Degree* (*D*): the number of the species’ links, within which we can distinguish between prey (*N_prey_*) and predators (*N_pred_*).
`o` *Betweenness centrality* (*BC*): it indicates how frequently node *i* lies on the shortest path between pairs of nodes *j* and *k*, thereby portraying its role as an “intermediary” in the network.
`o` *Closeness centrality* (*CC*): it measures how “close” a given node is to other nodes in the network, i.e., the average number of steps it takes to reach other nodes. The higher the value, the more directly the deletion of this group affects the majority of other groups.

- *Keystoneness* (*K*) (Jordán et al., 1999): while the degree of a node simply represents the number of nodes directly connected to it, the keystone index provides additional information on how these neighbors are interconnected with their own neighbors. This index quantifies vertical relationships such as trophic cascades but does not take into account horizontal interactions like apparent competition.

Another important factor to take into account is the similarity among the invaders concerning their interactions with prey and predators. The greater the shared predators and prey among them, the higher their perceived similarity. To quantify the extent of link overlap (*L*), we employed the Jaccard index as the index of similarity (Martinez, 1993):

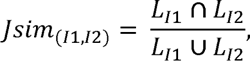

where *L_I1_* and *L_I2_* refer to the links belonging to *I1* and *I2*, respectively. The nature of the ecological relationship between invaders can also influence both the outcome of invasion and the degree of additivity (White et al., 2006; Preston et al., 2012). We defined the following relationship categories (and combinations of them, as shown in Fig 2.):

- Direct, *one-step trophic link* between the invaders:

`o` predator-prey relationship (*PP*);
`o` intra-guild predators (*IGPred*);
`o` intra-guild preys (*IGPrey*);
`o` their combinations (*IGPred*+*IGPrey*)

**Fig. 2.**
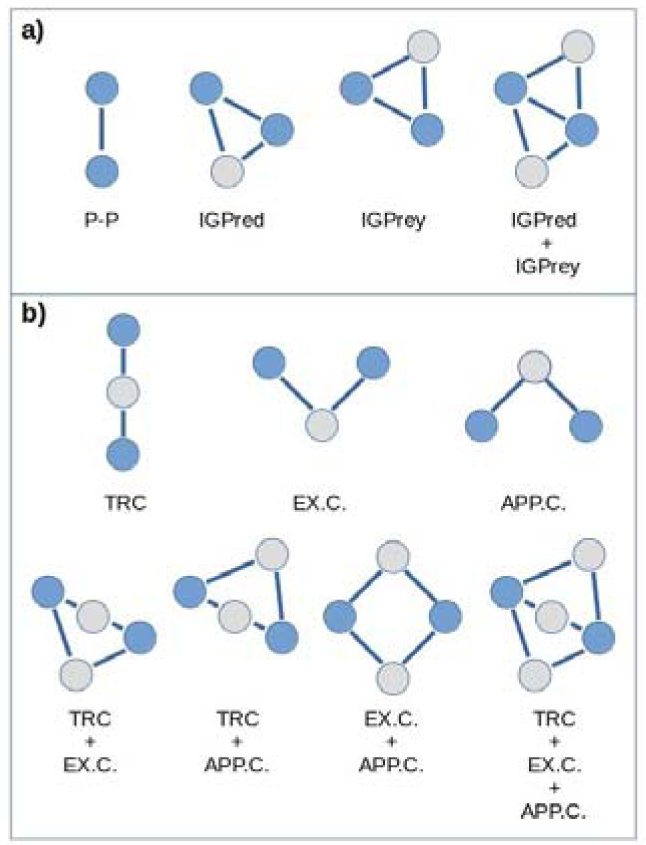
Investigated relationhip-categories between the two invaders (blue circles) a) 1-step distance links with feeding relationship (PP: predator-prey; IGPred, IGPrey: intraguild predator, intraguild prey; IGPred+IGPrey: both together); b) 2-step distance links (TRC: trophic cascade; EXC and APPC: exploitative and apparent competition; and their combinations)

- *Two-step links*:

`o` trophic cascade (*TRC*);
`o` exploitative competition (*EXC*);
`o` apparent competition (*APPC*);
`o` their combinations (*TRC+EXC*, *TRC*+*APPC*, *EXC*+*APPC*, *TRC*+*EXC*+*APPC*)

- *Distant links*: they are more than two-steps apart from each other.

## 3. RESULTS

### 3.1 Frequencies of different invasion outputs

One obvious way of determining the outcome of an invasion event is to record whether an invader is present in steady state. In this study, we tracked and compared the outcomes of two scenarios where the two invaders can be appeared in the system either separately or together.

Looking at success from a presence/absence perspective, we can distinguish 10 different outcomes, as shown in Fig. 3. along with their frequencies among all (121000) invasion events examined. These so-called invasion outputs (hereafter *IOP*), ranked according to their frequency from the most frequent to the least frequent one as follows:

A. Both of the invaders are successful alone and also together.
B. Only one is successful alone, and the same invader is successful in the simultaneous invasion, too.
C. Both are successful alone, but one is unsuccessful in the presence of the other.
D. One of them is unsuccessful alone, but both are successful when invading together.
E. No one is successful, neither alone nor together.
F. One of them is successful alone, but both are unsuccessful when invading together.
G. Both are unsuccessful alone but successful together.
H. Both are unsuccessful alone, but one is successful when the other is present.
I. Both are successful alone but unsuccessful together.
J. They “replace” each other: the one is successful alone, but in the simultaneous case, the other survives only.

**Fig. 3.**
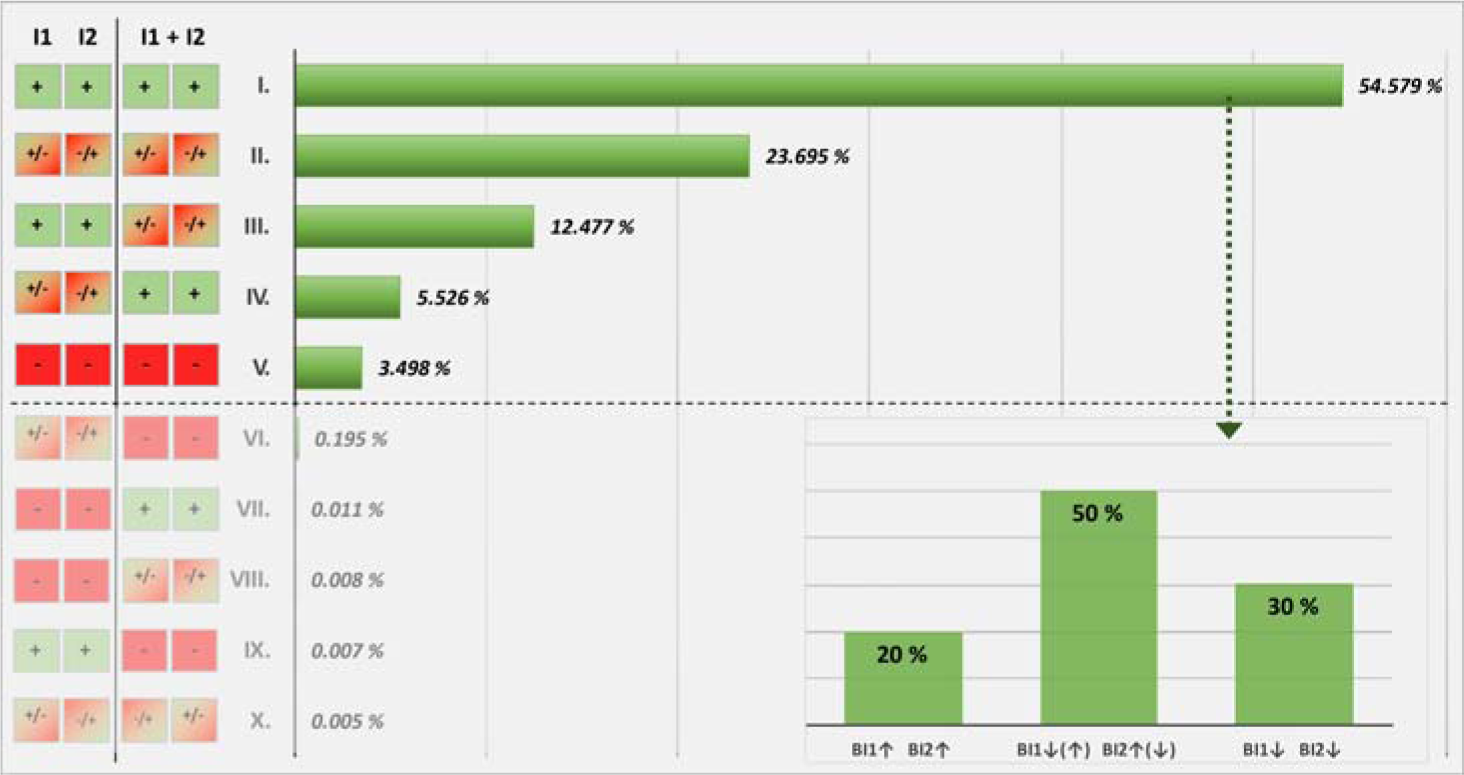
The potential outcomes of separate and joint invasions. The separate invasions are represented in the first two columns, while the outcomes of their simultaneous invasion are shown in the second two columns of squares. Successful invasions are depicted with green boxes and a “+” sign, while unsuccessful invasions are represented by red boxes and a “-” sign. Squares with mixed colours indicates scenarios where only one of the invaders succeeds. Due to the negligible frequency of cases VI-X (shaded), we focus on the first five cases (I-V.) for further analyses. The barplot in the bottom right corner shows the distribution of different categories of biomass changes within the always successful invasion output (IOP-I; ↑: increasing biomass; ↓: decreasing biomass).

Since the frequency of the cases VI-X is negligible (see Fig. 3), we focus only on the first five most frequent cases for the further analyses. IOP-I is a far more common case than any other (54.6% vs 23.7%, 12.5%, 5.5% and 3.5%). This high probability of success of separate and combined invasions is probably due to the well-known high-level stability of the network and the model settings (allometry, using the average Hill exponent), which remain largely unaffected following the invasion of one or two new species.

Since invasions are successful both in separate and joint scenario in the IOP-I case, it is possible to compare the equilibrium biomass of invaders between the two scenarios, i.e., to examine whether the biomass of each invader increases or decreases in the presence of the other compared to the separated case. This provides another aspect of the interaction between the invaders and an opportunity to examine the impact of different influencing factors. As shown in Fig. 3, in about 20% of cases the biomass of both invaders increases when they co-invade comparing to the case when they invade separately, in about 50% of cases, the biomass of one increases while that of the other decreases in co-invasion, and in 30% of cases, the two invaders mutually inhibit each other’s growth, leading to a lower equilibrium biomass compared to when they invade separately.

### 3.2 Distribution of the invaders’ ecological relationship-categories

As mentioned in section 2.4, ecological relationship between two invaders is divided into three major groups: i) direct trophic link, where the two invaders are 1 step apart, ii) not a direct, but a contact two steps away connection and iii) a distance of more than two steps. As Fig. 4 shows, in 73% of the results (IOP I-V only), they are at most two steps apart, in 27% of the results, the distance between them is more than two steps. The latter is treated as a single category in the following analysis, while the first two are subdivided into the already defined sub-categories (see Fig. 2. and Fig 4.).

**Fig. 4.**
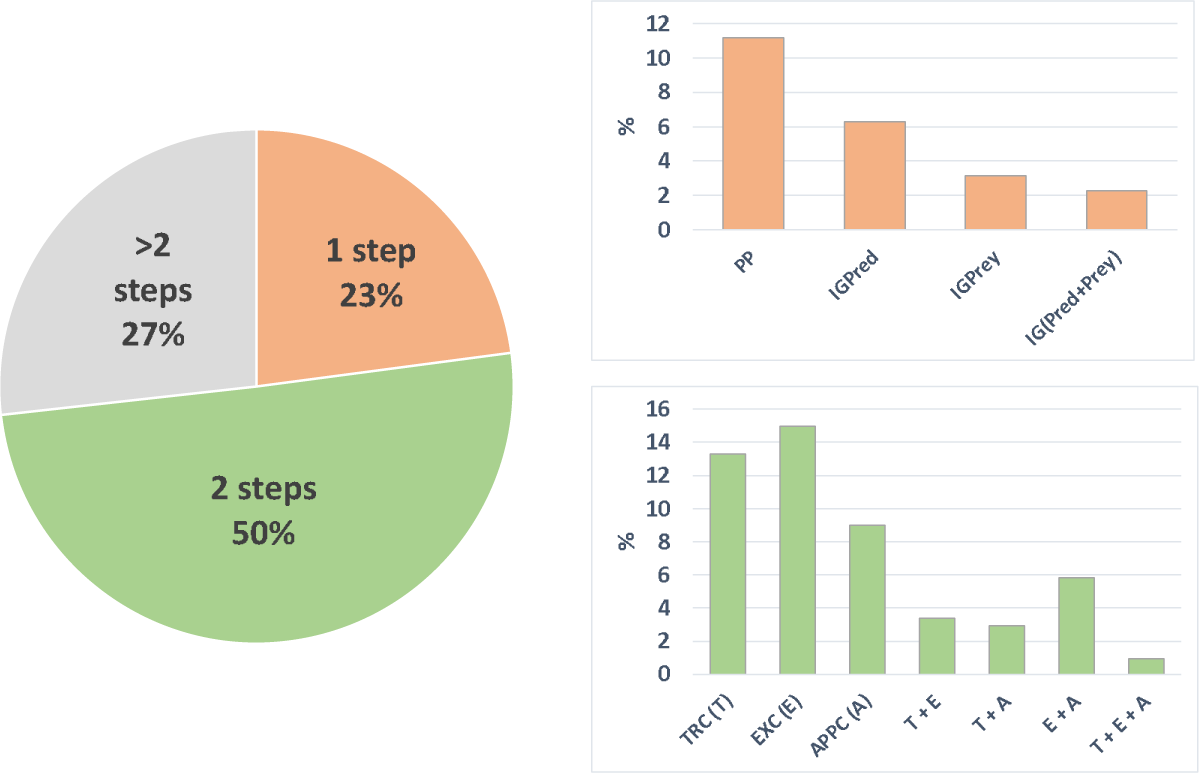
Frequency of invasion events as a function of the distance of the two invaders in the network (left), and the frequency of invasion events in each subcategory of the 1- and 2-step distance groups (right).

There is some similarity in the distributions of these sub-categories: the frequencies of category-combinations (e.g., *IG(Pred+Prey)* or *TRC*+*APPC*) are lower than that of the simple predator-prey (*PP*) or trophic cascade (*TRC*), and in both cases, it can be observed that the invaders share a common prey more often than a common predator (compare IGPred with IGPrey or EXC with APPC in Fig. 4.).

### 3.3 Is the nature of the ecological relationship between the two invaders related to the outcome of the invasion scenarios?

The effect of the type of ecological relationship and the positions of the two invaders in the food web was tested using chi-square independence tests with Cramer’s V effect size metric. (Table S3. in Supplement 2). For each (sub)category, we applied a chi-squared goodness-of-fit test with Cohen’s ω effect size metric to examine the difference in the frequency of outputs between that (sub)category and the full sample (Fig 3. and Fig 5.). This was carried out for different invasion outcomes (IOP I-V) and for different directional changes in biomass that were achieved in the course of a successful invasion (IOP-I). In the following, we only report here the values of the effect sizes; for more details on the statistics, see the Table S2. in the Supplement 2.

**Figure 5.**
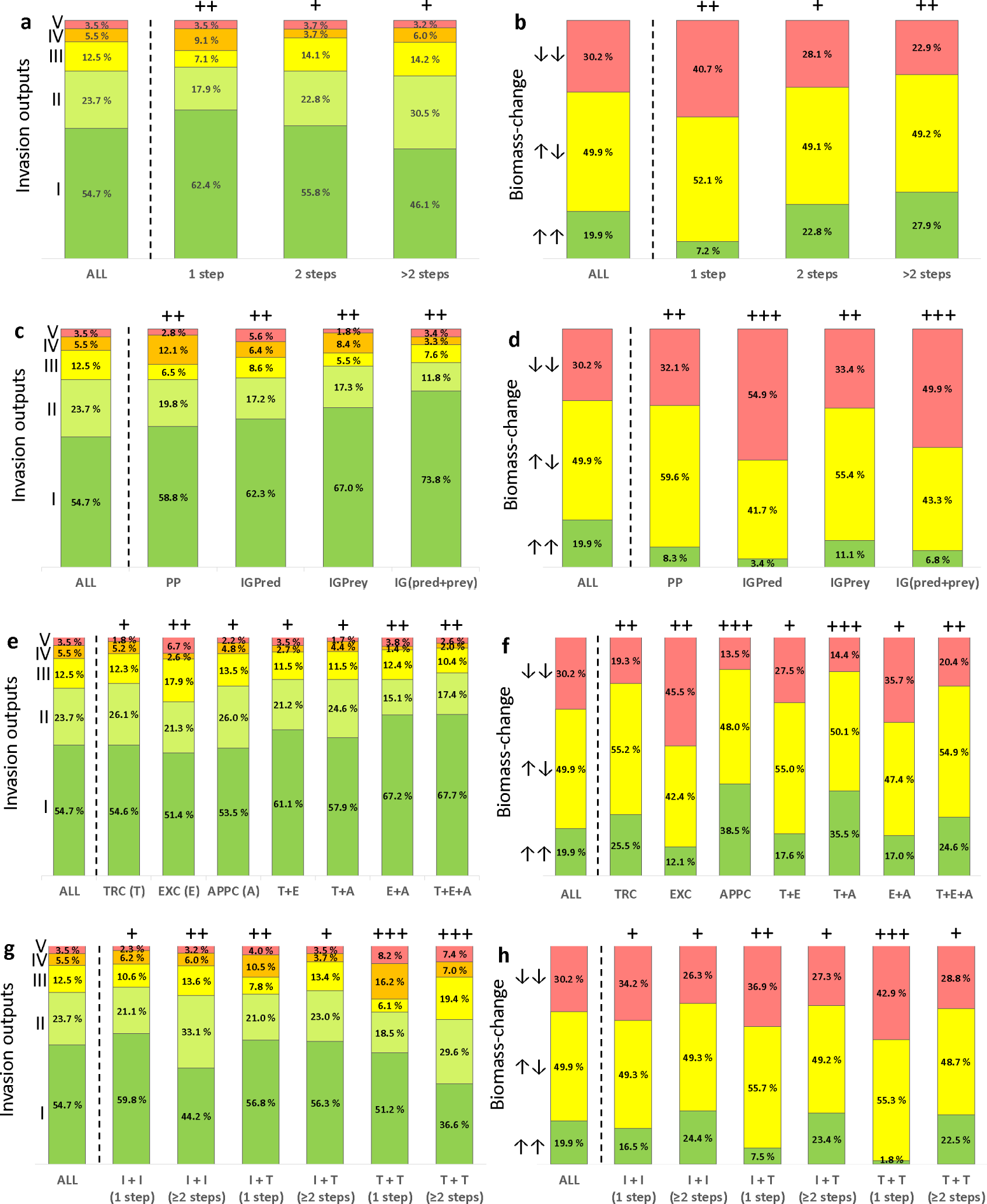
Frequency distributions of the different invasion outputs (IOP I-V, left panels) and the biomass-changes of the invaders in various directions distinguished by arrows (right panels) are shown within the (sub-)categories of the invaders’ ecological relationships (a-f), and, in function of their intermediate (I) or top (T) positions (g-h). The first column shows the frequency distribution across all outcomes (ALL), while the symbols above the rest of the columns indicate whether the effect size (Cohen’s ω) is small (+), moderate (++) or large (+++), indicating the magnitude of the deviation from the full sample.

The topological distance between the two invaders has a small effect on the frequency of both the different invasion outcomes (Cramer’s V = 0.12, Fig. 5a) and the on the direction of biomass change (Cramer’s V = 0.15, Fig. 5b). However, clear trends can be observed: in the case of a closer, 1-step predator-prey relationship the goodness-of-fit and post-hoc tests show significantly higher proportions of always successful invasions (IOP-I, 62.4% vs 54.7%), and also for those where only one invader is successful in the separate case, but both are successful together (IOP-IV, 9.1% vs 5.5%) than in cases where the topological distance between the invaders is larger (Cohen’s ω = 0.27). In contrast the proportions of cases where only one invader is successful (IOP-II) and where one is unsuccessful when invading together despite being successful in the separate cases (IOP-III) are increased comparing to the cases where topological distance between the invaders is 1 (30.5% vs. 17.9% and 14.2%, vs 7.1% respectively; Cohen’s ω = 0.19). Furthermore, if we consider the distribution of the biomass changes of different directions (Fig. 5b), we see that one-step relationship between the invaders greatly increases the likelihood that the equilibrium biomass of both invaders decreases (40.7%, vs 30.2%, Cohen’s ω = 0.34), and the more distant the invaders are from each other in the network, the more rarely have this negative effect on each other.

Studying the subcategories within the 1-step distant relationships (Fig. 5c), it is also true that in spite of the weak association (Cramer’s V = 0.09), the closer the relationship is, i.e. the more links there are between them, the more likely it is that invaders that are successful individually will also be successful together (increasing proportion of IOP-I from PP (59%) to IG(pred+prey) (74%)). However, we observe a difference between intraguild predator (*IGPred,* Cohen’s ω = 0.23) and prey (*IGPrey,* Cohen’s ω = 0.33) links. In the former case, there is a significantly higher proportion of invasions that always fail (IOP-V, 5.6% vs 1.8%) and a relatively higher proportion of IOP-III (one of the invaders that succeed separately fails in the simultaneous case) (8.6% vs 5.5%). In the latter case, it is precisely the proportion of IOP-V that decreases significantly (1.8%), while that of IOP-IV increases (one fails separately, but they succeed together) (8.4% vs 6.4%). However, when considering the directions of biomass changes (Fig. 5d), only the *IGPred* relationship has a strong effect on the distribution (Cohen’s ω = 0.59), not only alone, but also in combination with a common predator (*IG(Prey+Pred),* Cohen’s ω = 0.47). In such a case, the probability that the equilibrium biomass of both is reduced during the joint invasion is significantly increased compared to the separate case. This effect cannot be compensated even if they also share a predator (54.9% and 49.9% vs 30.2%).

The connections between the 2-step distance subcategories (and their combinations) and IOP-distributions (Fig. 5e) is even weaker (Cramer’s V = 0.09). Nevertheless, it can be concluded that, as before, an increase in the tightness of the relationship (the number of links) increases the probability that invaders will be successful not only alone but also together. The frequencies for *TRC* and *APPC* are not remarkably different for any of the outcomes, neither from each other nor from the frequency distribution of the whole sample (Cohen’s ω = 0.1), the *EXC* (shared prey) relationship significantly increases the odds of complete failure (IOP-V, 6.7% vs 3.5%) and that of one of the two invaders succeeding alone and the other being extirpated (IOP-III, 17.9% vs 12.5%; Cohen’s ω = 0.27). However, if the invaders share not only a common prey but also a common predator (*EXC*+*APPC*) or are in a trophic cascade relationship as well (*TRC*+*EXC* or *TRC*+*EXC*+*APPC*), this effect is overridden, and the probability of success (IOP-I) increases (61.1% or 67.7% vs 54.7%). In respect to biomass-changes (Fig. 5f), the most pronounced effect is that of apparent competition (*APPC*), which significantly increases the probability that the two invaders will have a positive effect on each other, i.e., the equilibrium biomass of both will increase when they occur together in the system (38.5% vs 19.9%, Cohen’s ω = 0.52). A smaller but similar directional effect is obtained when there is a trophic cascade relationship between them (25.5% vs 19.9%, Cohen’s ω = 0.25). Exploitative competition, on the other hand, has the opposite effect: as with intraguild predation, the probability of biomass decline increases significantly for both invaders when prey is shared (45.5% vs 30.2%, Cohen’s ω = 0.35). Only its combination with both *TRC* and *APPC* can substantially compensate for the effect of the *EXC* relationship (27.5% or 20.4% vs 45.5%).

We also examined whether the frequency of different outcomes was affected by the invaders occupied intermediate (I) or top (T) predator positions (Fig. 5 g-h). Although we found a weak overall relationship between the two variables (Cramer’s V = 0.1), the post-hoc test showed that the probability of successful invasion for both scenarios was higher when at least one invader occupied an intermediate position or became intermediate when its predator appeared. When both invaders occupy intermediate positions (I+I), the existence of a predator-prey relationship between them is the determining factor of the probability of success (59.8% vs 44.2%). When one invader occupies an intermediate position and the other is an apex predator (I+T), the predator-prey relationship does not increase the proportion of successful invasions in either scenario, but the difference between IOP-III and IOP-IV becomes apparent: the closer, predator-prey relationship increases the probability that an invader that is unsuccessful alone, survives in the presence of the other (10.5% vs 3.7%). In addition to the fact that the top position increases the invasion failure rate, when one top position invader consumes the other top position invader (which thereby shifts one of them to the intermediate position) during the joint invasion, they are more likely to help each other than when there is no predator-prey relationship between them (IOP-IV, 16.2% vs 7%). If both invaders occupy top positions, the likelihood of invasion failure increases significantly, particularly when there is no predator-prey relationship between them (Cohen’s ω = 0.4). In such cases, not only is the probability of total failure (IOP-V) considerably higher (7.4% vs the overall 3.5%), but also the likelihood of either of them going extinct when they jointly invade, irrespective of whether its invasion was initially successful (IOP-III, 19.4% vs the overall 12.5%) or not (IOP-II, 29.6% vs the overall 23.7).

All in all, the key distinction lies not primarily in their position, but rather in the presence or absence of a predator-prey relationship, and it is true in the context of the biomass-change categories, too (Fig. 5h). As already demonstrated in Fig. 5b 1-step connections typically exhibit a lower likelihood of boosting the biomass of both invaders compared to more distant relationships. While the position in itself doesn’t exert a significant impact in the context of distant links, in the case of a predator-prey link, occupying the top position increases the likelihood of a biomass decrease for at least one invader (55.3% vs the overall 49.9%), and often both of the invaders (42.9% vs the overall 30.2%) with a large effect size (Cohen’s ω = 0.47).

### 3.4 Which invader survives in case of partially successful joint invasions?

In case of IOP-II and -III one of the invaders faces failure when invading together. Furthermore, in the case of IOP-I, the biomass of one invader increases while the other’s decreases in half of the cases (Fig. 3). These observations raise the question of whether we can determine, based on their ecological interactions or topological indices, which invader is more likely to survive or increase in biomass.

When it comes to the two competition categories (exploitative or apparent), there is no clear topological hierarchy (see Fig. 2b) that inherently indicates which invader can be favoured. However, for the remaining categories (including combinations with competition), distinctions can be drawn, such as between prey and predator or, in the case of trophic cascades, between lower and higher-position invaders. Note that, as mentioned in section 2.3, invaders were designated as *I1* and *I2*, and although they were introduced to the resident community very close in time, *I2* was always placed “*above” I1*. This means that the success of *I1* always implies the success of the lower or prey position. The results of the comparisons of the probabilities of *I1*’s or *I2*’s success as a function of the different ecological relationships are shown in the panels of Fig. 6, the details of the statistical analysis are summarized in the Table S6.

**Figure 6.**
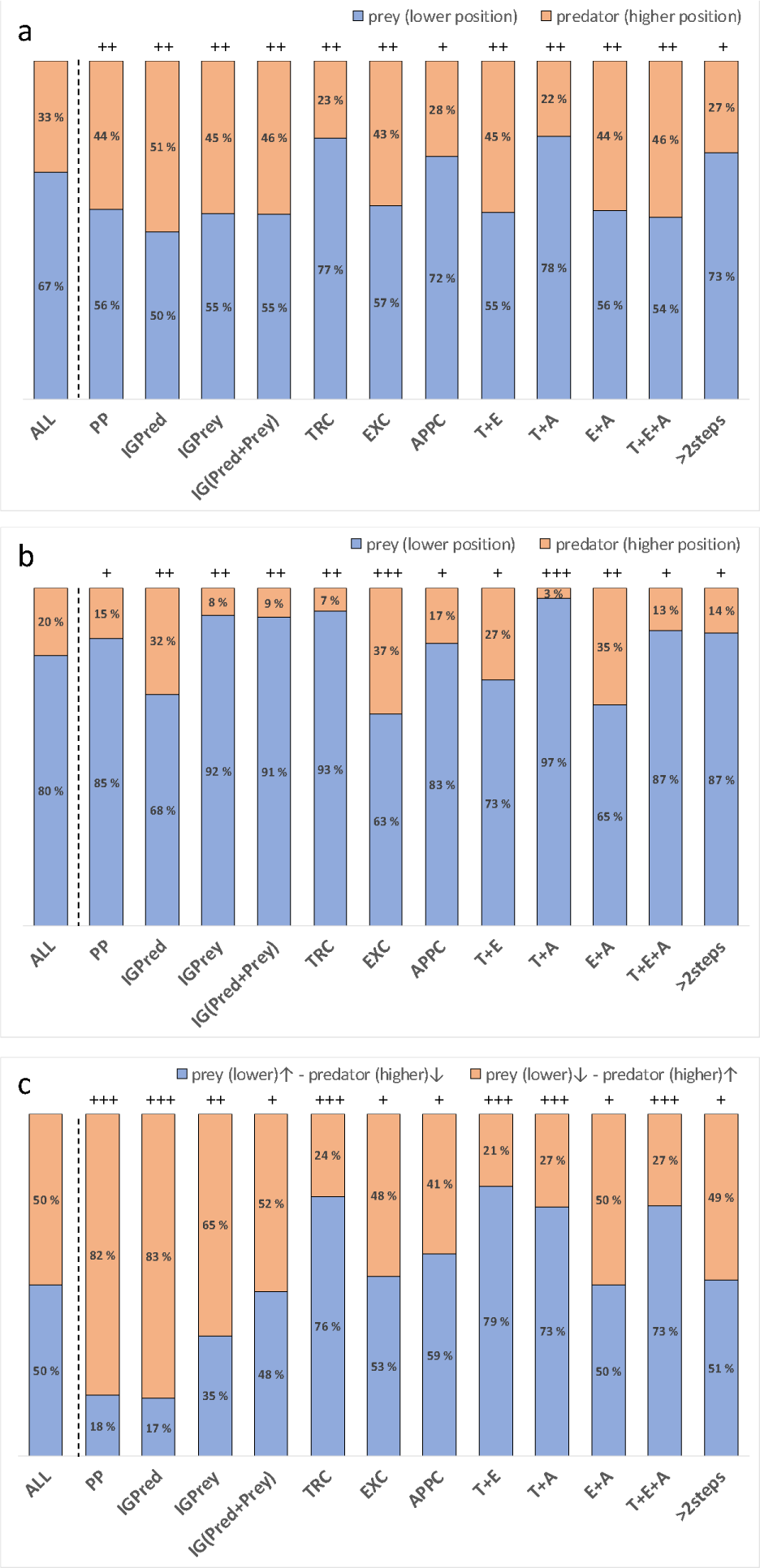
Frequencies of the success of the invader in prey/ “lower” position (blue) or in predator/”higher” position (orange) in case of IOP-II (a), IOP-III (b) and the direction of biomass-changes denoted by arrows found these positions in case of IOP-I (c). The first column shows the frequency distribution of the success of the invaders of different role (ALL), while the symbols above the rest of the columns indicate whether the effect size (Cohen’s ω) is small (+), moderate (++) or large (+++), indicating the magnitude of the deviation from the overall frequency distribution within the output case in question.

In the case of IOP-II (one invader fails in both the separate and joint cases), the overall probability of *I1* in the lower (prey) position succeeding is 67%, while the probability of *I2* (predator or “higher” positional invader) succeeding is 33% (Fig. 6a). There is a connection between the topological arrangement of invaders and the probability of predator’s or prey’s success, however, this relationship is relatively weak (Cramer’s V = 0.2). In the goodness-of-fit test, we examine how the frequency of the success/failure varies across the ecological categories compared to this observed frequency. We observe that in each of the 1-step predator-prey relationships, there is an increase in the likelihood of success for *I2*. The largest deviation from the observed overall frequencies occurs due to intraguild predation (*IGPred,* 51% vs 33%, Cohen’s ω = 0.37), wherein they even share a common prey, consequently further elevating the probability of success in the predator position. In contrast, the trophic cascade (*TRC*) relationship further increases the success rate of the lower position *I1* (77% vs 67 %, Cohen’s ω = 0.21).

We would expect that in competition categories where we cannot differentiate between top and bottom positions, there wouldn’t be a significant disparity in the success rates of the two invaders. This holds true for exploitative competition, which raises the success rate of *I2* from 33% to 43%. Interestingly, however, apparent competition does not tend to equalise the success/failure rates; instead, it further increases the probability of success for I1 (72% vs 67%). Although the effect size in this case is small (Cohen’s ω = 0.1), when combined with the trophic cascade, it leads to a further increase in *I1*’s success rate (78% vs 67%, Cohen’s ω = 0.23). On the other hand, exploitative competition (EXC) counteracts the effects of both trophic cascade (TRC+EXC, Cohen’s ω = 0.25) and apparent competition (APPC+EXC, Cohen’s ω = 0.24), increasing the success rate of the higher-positioned *I2* (from 30% to 45% and 44%, respectively).

In the case of IOP-III (Fig. 6b) unlike the prior scenario, both invaders achieve success individually, but one of them goes extinct when they invade the resident food web together. Notably, the likelihood of the prey or the lower-ranking invader (*I1*) sustaining success is notably higher (80%). Remarkably, unlike the previous situation, only the intraguild predator (*IGPred*) interaction enhances this probability within the 1-step categories by increasing the predator’s (*I2*) survival prospects (32% vs 20%, Cohen’s ω = 0.29). However, if a common predator is present (*IGPrey*), it further diminishes the chances of *I2* surviving (8% vs 20%, Cohen’s ω = 0.28). The connection with EXC remains consistent with the previous scenario, where it notably narrows the gap between the survival probabilities of the two invaders with a large effect size (57% vs 47% for *I1* and *I2*, respectively, Cohen’s ω = 0.44). However, the same cannot be said for APPC (72% vs 28% for *I1* and *I2*, respectively, Cohen’s ω = 0.08), as well as the TRC relationship significantly increase the success rate of the invader in the lower position (77% vs the overall 67%, Cohen’s ω = 0.32).

In IOP-I, in half of the cases, the equilibrium biomass of one invader increases while that of the other decreases in case of joint invasion (see Fig. 2), as compared to what each achieved when present in the system individually. The probability of the biomass of either I1 or I2 increasing is precisely 50% (Fig. 6c). Therefore, we can assess the variation within each category with respect to this ratio. If they form a predator-prey relationship during the joint invasion, thus, *I2* consumes *I1*, it’s more probable for the predator to achieve a higher equilibrium biomass compared to when it exists alone in the system (PP: 82% vs 50%, Cohen’s ω = 0.64; IGPred: 83% vs 50%, Cohen’s ω = 0.66). The influence of a shared predator (IGPrey and IG(Pred+Prey) diminishes this advantage significantly but doesn’t shift it toward the prey (65% vs 50%, Cohen’s ω = 0.3 and 52% vs 50%, Cohen’s ω = 0.04, respectively). However, in the category of trophic cascade, the opposite holds true, with the lower-positioned invader being more likely to increase its equilibrium biomass (76% vs 50%, Cohen’s ω = 0.51). This trend remains consistent even when trophic cascade effects combine with any type of competition (TRC+EXC: 79% vs 50%, Cohen’s ω = 0.58; TRC+APPC: 72% vs 50%, Cohen’s ω = 0.46). It’s worth noting that neither any type of competition nor links with a distance greater than 2 steps have a substantial impact on either increasing or decreasing biomass. In the case of apparent competition, there’s also a slight favor towards biomass increase of *I1* (59% vs 50%), but the effect size is small (Cohen’s ω = 0.18).

### 3.5 Connection between the invaders’ network indices or similarity and the invasion outcomes

The correlation of the topological position of the two invaders can also captured by the difference (*d-’IND’*) and average of the network indices (*a-’IND’*) expressing them. We also investigate whether there is an effect of the invaders’ similarity (*Jsim*) on the invasion outcome. We tested whether there is a significant difference between the differences and averages of these indices for each invasion outcome and, if so, how strong it is (η2 effect size). The most relevant results are presented in the panels of Fig. 7., the details for all investigated indices and the statistics can be found in in the Supplement 2 (Table S4 and S5). In general, it is true for all indices that their differences have a much smaller effect than their averages (see Table S4), i.e. the latter tend to determine the differences between the invasion outputs. However, this effect is small (η2 < 0.06) or negligible (η2 < 0.01) for almost all network indices, the invaders’ trophic level has the strongest influencing effect (η2 = 0.09, see Table S4.). Here we highlight results for indices for which significant relationship has been identified; results for other indices are shown in the Supplement 1 (Fig S1-S4).

**Fig. 7.**
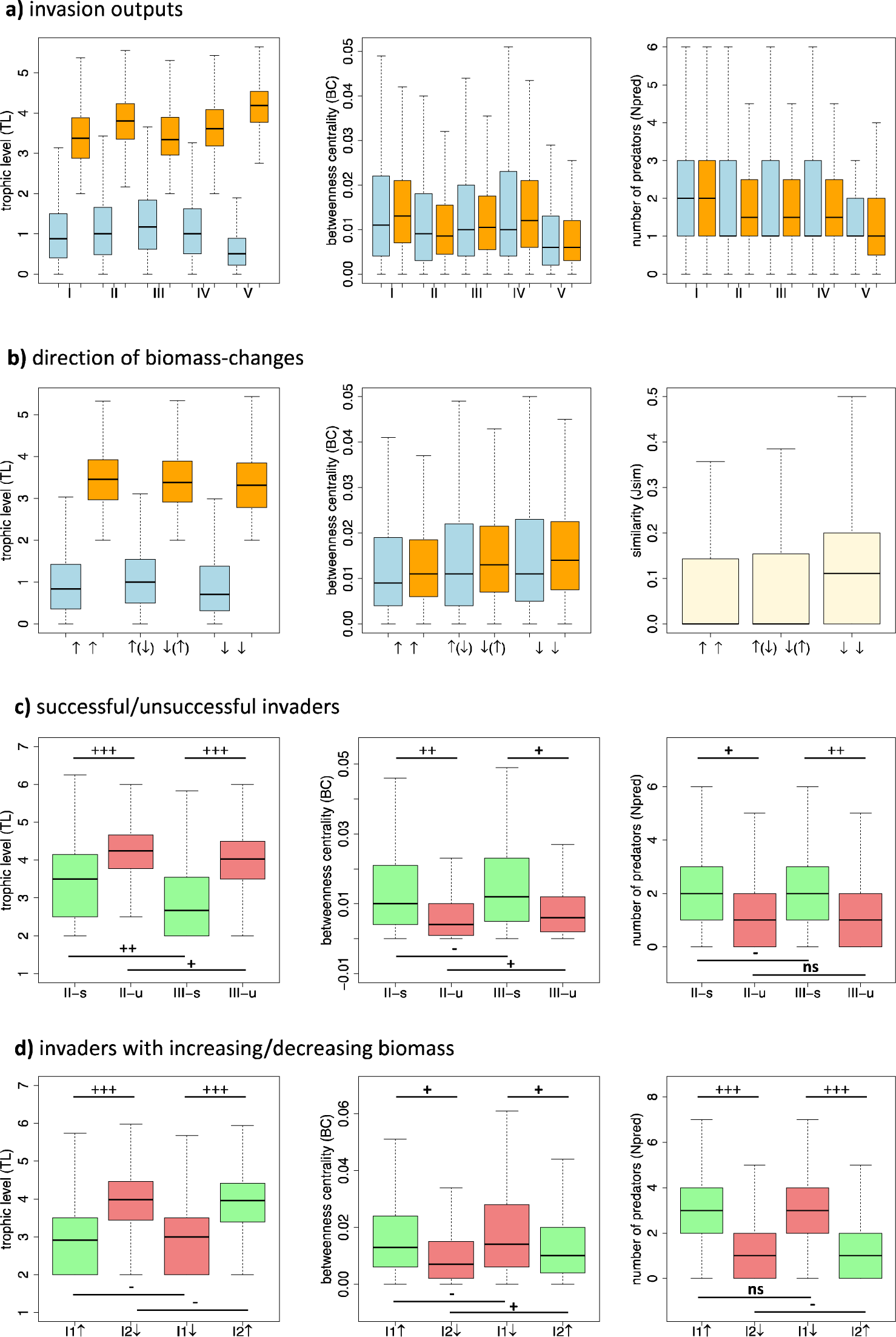
The differences (blue) and the averages (orange) of the invaders’ highlighted positional network indices and similarity values (beige) in case of **a**) the different invasion outputs (IOP) and **b**) of invaders for different directions of changes in their biomass in case of IOP-I (↑↑: both increase; ↑(↓) ↓ (↑): one of them increases, the other decreases; ↓↓: both decrease). **c**) Distributions of the successful (s, green) and unsuccessful (u, red) invaders’ highlighted network indices in case of IOP-II and IOP-III. **d**) Distributions of the topological indices specific to invaders with increasing (↑, green) and decreasing (↓, red) biomass in case of IOP-I. The lines above and below the boxplots represent the cases being compared. The strength of the relationship is indicated by Cohen’s d effect size, which is categorized as non-significant (ns), negligible (-), small (+), moderate (++), or large (+++).

We found the largest differences between the IOP-categories in terms of the invaders’ trophic level (*TL*). IOP-V, i.e. the case of complete failure, is the most different from the other outcomes, with significantly higher average *TL*s of the invaders (Fig. 7a). IOP-II (one invader fails) also has higher a-TL with a medium effect size compared to IOP-I (Cohen’s d = 0.64) and IOP-III (Cohen’s d = 0.56). IOP-III and -IV are not considerably different, with small (|Cohen’s d| < 0.5) or negligible (|Cohen’s d| < 0.02) effect sizes. In the case of betweenness centrality (*BC*), IOP-V shows a moderate degree of divergence with respect to the others (except for IOP-II, where one invader also fails; Cohen’s d = -0.29). Furthermore, in this outcome, the invaders have less predators in average (a-*Npred*) (Cohen’s d = -0.61) than that of in the case of IOP-I. This is consistent with a higher a-*TL* where species are more likely to be top predators than at lower levels of the food web.

However, the direction of the invaders’ biomass changes investigated for IOP-I are not or only slightly influenced by their topological indices (see Fig. 7b, Fig S2 and Table S4 in Supplemet 1, 2.), even by their trophic level. The overall effect size is negligible everywhere (η^2^ < 0.01), the only small difference is revealed for betweenness centrality and similarity: slightly higher a-*BC (*Cohen’s d = 0.26) and *Jsim (*Cohen’s d = 0.29) are more likely to decrease the equilibrium biomass of both invaders.

### 3.6 Connection between the invaders’ network indices and their success/failure

We also investigated whether distinctions exist between the topological network indices of successful and unsuccessful invaders in cases IOP-II and IOP-III. The results of the statistics are summarized in Table S7. Our visual representations (Fig 7c, Fig. S3) clearly show that the most substantial difference is observed again within the trophic level of the two invaders. Successful species consistently exhibit significantly lower TL-values compared to unsuccessful invaders, irrespective of outcome type (IOP-II or -III, Cohen’s d = -0.99 and -1.43, respectively). In contrast, the reverse is true for betweenness centrality. For both outputs, higher BC-values are associated with a higher probability of invasive species’ success, albeit with less significant impact (Cohen’s d = 0.6 and 0.48 for IOP-II and -III, respectively). Successful invaders have a higher average number of predators in both outcomes (Cohen’s d = 0.45 and 0.53 for IOP-II and -III, respectively).

Interestingly, we do not see any difference in the topological network indices for biomass increase or decrease in case of IOP-I (Fig. 7d. Fig. S4 and Table S7 in the Supplement 1, 2). Whether joint invasion increases its biomass or not, the trophic level of *I2* is always higher than that of *I1*. The same is true for the number of invaders’ predators. It is true for all the indices investigated that whether the average of the groups analysed is slightly or significantly different, the differences are not correlated with an increase or decrease in biomass, but are consistently observed between the two invasive species. This suggests that network indices in isolation do not influence the increase or decrease in biomass of invasive species following a simultaneous invasion.

## 4. DISCUSSION

We assessed the invasion success and interactions of two invasive species in model food webs, utilizing two approaches: one based on their presence or absence, and the other based on changes in biomass, whether it increased or decreased in the presence of each other compared to the separate situation. These two aspects produce results that show both similarities and differences. Our research is structured around four questions. In the following, we summarise the answers to the questions based on our results.

### How do the invaders interact when they appear in a resident community together?

In the majority of cases, both invaders were able to spread in the system, not only individually but also in the presence of the other (IOP1 scenario). This is due to the allometry-based model, which lends a high degree of stability to the network and thereby to the invasive species. However, this does not imply the absence of mutual influence, as changes in their equilibrium biomass can be traced to determine which is more successful. From this perspective, in 80% of our findings, one of the two species has a lower equilibrium biomass when in the presence of the other. Only in 20% of cases can it be concluded that they mutually benefit each other from biomass increase aspect. The possible reason for this is that in our food web model, mutualism can only occur through indirect cascading effects, while direct effects are clearly negative (competition, predation).

### Do simultaneous invasion outcomes depend on direct and indirect ecological interactions between invaders?

The closer the relationship between the two invaders topologicaly, the more likely they are to succeed both separately and together, with a corresponding decrease in the frequency of at least one of them going extinct in the presence of the other. However, the opposite result is obtained for biomass changes, with distant relationships resulting in a larger proportion of biomass increases for both invasives compared to the separated case (see Fig. 5).

An important distinction arises when considering if invading species share a common resource or predator. If they share a common prey, either through predator-prey relationships or competition for resources, it increases the likelihood of both invaders failing in either scenario. If they fail independently, it also increases the likelihood that neither would succeed even after co-invasion (IOP-V). On the other hand, if these invaders share a natural enemy, either through intra-guild predation or apparent competition, this increases the likelihood that an otherwise unsuccessful invader will survive in the presence of the other (IOP-IV). In such cases, both invaders may even experience an increase in biomass compared to separate invasions.

The outcome of invasion is influenced by whether the two invasive species occupy a top or intermediate position in the network. Invading in the top position, which typically corresponds to a higher trophic level, there is a higher probability of invasion failure. However, another key factor is the presence or absence of a predator-prey relationship. Rather surprisingly, if there is a direct feeding link between the invaders, this increases the success rate of both. Conversely, if they are far apart, it’s more likely that at least one will not survive. This result is in contrast to the biomass change approach, where a distant relationship is more likely to result in an increase in biomass for both invaders for joint invasion, while the presence of a direct predator-prey relationship will decrease the biomass of at least one, and often both, of them.

### How do different topological indices have an impact on the invasion outcome of invaders within the food web?

Assessing the impact of network indices, our findings indicate that they exert a significant influence only in the presence/absence approach. In this context, it becomes evident that the trophic level of invaders plays a crucial role (see Fig. 7). Consistent with prior research (Romanuk et al., 2009), a higher trophic level tends to increase the likelihood of unsuccessful invasion for at least one invader. Regarding the other indices, we observe minor or negligible differences in both the average values and the disparities between these indices. It is noteworthy that among the different outcomes, the scenario where both invasive species fail stands out more clearly from the rest. This is not only in terms of having a higher average trophic level but also lower average betweenness centrality, fewer predators, and higher average similarity. However, these distinctions are closely tied to the fact that these biases in the indices are all associated with a higher trophic level, mostly at the top position in the food web.

We have found that there is no meaningful relationship between the direction of biomass-change and the topological indices of the two invasive species, even when their trophic level is taken into account. We do observe a slight distinction in terms of betweenness centrality and average similarity: the biomass of the invaders is on average more likely to decrease for higher betweenness centrality and similarity, compared to when they exist separately. However, this effect is relatively small. Therefore, it can be concluded that the biomass-change is not primarily correlated with the various topological indices, but rather by the nature of the direct or indirect relationships between the two invasive species.

### If only one invader is successful, which one is it? Furthermore, which positions or topological indices indicate its’ success?

Our findings are further supported by the comparison of the likelyhood of success/ increasing biomass according to the invaders’ different positions (prey(”lower”) / predator(”higher”)) and their topological indices. When one invader’s biomass increased while the other’s decreased during simultaneous invasion, no significant differences were observed in the indices, even at their trophic level. It became evident that the direct and indirect interactions between these species played a much more influential role in explaining biomass change by joint invasion: in cases involving a predator-prey relationship, the predator might reduce the biomass of the prey, whereas in scenarios featuring a trophic cascade relationship, the invader at the lower trophic level responded with biomass increase while the one at the upper level experienced biomass decrease, comparing to their equilibrium biomass after invading the resident food web separately. In situations where there was no clear topological hierarchy (e.g. competition), the likelihood of both species experiencing an increase or decrease in biomass because of the joint invasion was nearly equal. This ratio is altered when there is a predator-prey relationship or a trophic cascade, with the predator being favoured in the first case and the invader being in a lower position in the second (Fig. 6).

If one of the two invading species goes extinct while the other successfully survives during their joint invasion, significant differences emerge in several ecological indices. In the majority of cases, the successful species tends to occupy lower trophic levels and exhibits higher betweenness centrality compared to the unsuccessful one. There is no significant difference in the number of prey species associated with them, but it’s noteworthy that the unsuccessful invaders typically have fewer predators (Fig. 7). When only one of the invaders is able to maintain a viable population within the ecosystem, in case of joint and separate invasion, the successful species is characterized by a higher degree number and a higher keystoneness index compared to the extinct species (Fig. S3).

The magnitude of the effect of the trophic level suggests that in most cases, an invader is more likely to survive in a lower position (e.g., within a trophic cascade) or as prey. This observation aligns with the idea that it is generally easier to establish oneself in an ecosystem as prey rather than as a predator (Jiabu and Li, 2023). However, when a shared prey is involved in addition to a predation relationship (*IGPred*), the predator’s chances of survival increase due to the additional energy provided by the shared prey and the potential for food switching. In cases of competition, the survival chances of invaders in under-over position are more evenly distributed. This also means, however, that invaders who fail individually are unlikely to coexist after joint invasion, but when two independently successful invaders are in competition, other factors such as propagule pressure or the timing of invasion may play a crucial role in invasion success (Lockwood et al., 2005).

### Outlooks and limitations

Examples of both simultaneous and sequential colonization can be found in real systems (Simberloff and Wilson, 1969; Carlton and Geller, 1993; Ruiz et al., 2000). The latter has led to the development of the concept of positive interactions between invaders, which was encapsulated in the Invasion Meltdown Hypothesis (Simberloff and Holle, 1999; Simberloff et al., 2013). It is a widely studied question whether a non-indigenous species, by invading, changes the community structure in a way that creates more favourable conditions for a subsequent invading species (Grosholz, 2005; Green et al., 2011; Zhang et al., 2020). This question is relevant not only in macroecological contexts but also in microbial communities (Herren, 2020) where the effects are often not solely trophic but also mediated by metabolite-mediated interactions, such as cross-feeding (Brunner and Chia, 2019). Most microbial invasion studies, similar to macroecological contexts, focus on two primary analytical directions (Mallon et al., 2015): the success of the invasive species and the resulting change in community composition (Kinnunen et al., 2016), as well as the the resilience of the community (De Roy et al., 2013). However, researchers are increasingly paying attention to cases in which, even though the invader cannot establish a permanent population in the system, short-term interactions with them can induce long-term changes in community functioning and composition, thereby facilitating subsequent invasions. Beyond these, however, increasing attention is being paid to cases where, although the invader is unable to establish a permanent population in the system, short-term interactions with them can induce lasting changes in community functioning and composition, thereby facilitating a subsequent invasion. Such a transient invasion may underlie the invasion outcome we found, but which has not been studied because of its rarity, where two invaders are not successful separately, but when they appear together (or in close succession), one can take advantage of conditions favoured by the other and spread (see IOP-VIII in Fig. 3). This phenomenon has been studied in laboratory settings (Amor et al., 2020), as its importance is not negligible, for example, in the human gut microbiome (David et al., 2014), but transient invasions have also been observed in freshwater phytoplankton communities (Buchberger and Stockenreiter, 2018).

An important question that arises is whether our understanding of multiple invasions and interactions among invasive species can be harnessed to manage the proliferation of aggressively spreading invasive species. One approach to biological control is to utilize species interactions as a means to control or eliminate invasive species (Hawkins and Cornell, 1999). Our results suggest that, although in the modelling framework we use, it is extremely rare for an invasive species to be displaced or ’replaced’ by another invader, this can happen through predator-prey relationships or exploitative competition. Russell et al., 2014 investigated the effects of competitive displacement between invasive rats, and found that functional similarity can lead to displacement through competition. However, the inherent unpredictability of indirect interactions is a major challenge in terms of their impact on invasions (White et al., 2014), so great caution is required in projects of this nature (Simberloff, 2012).

In our research, we employed an allometric model rooted in a bioenergetics perspective. This approach is of moderate complexity and provides a more accurate picture of species interactions than the Lotka-Volterra models commonly used in the study of species-rich systems. It simplifies the system by using allometry being common in nature, and ignores many processes to avoid over-parameterization and model complexity (Yodzis and Innes, 1992), thus using the allometric model has significantly enhanced our understanding of community functioning and stability and has found successful applications across various ecosystems, predominantly in aquatic systems, but also in terrestrial soil communities (Perkins et al., 2022). Nevertheless, a limitation of this method lies in its suitability for terrestrial ecosystems, where body mass is not always a reliable predictor of plant-consumptive interactions. Recent efforts have aimed at developing a bioenergetics framework that integrates these processes (Valdovinos et al., 2023), potentially rendering it suitable for studies like ours in the context of terrestrial invasions. Further, like the Lotka-Volterra model, this approach does not directly address the problem that species at different life-stages can be in different position within the food web, which would be interesting to study in the future.

Another limitation of our study is that, for the sake of tractability, we heavily focused on the properties and interactions of the two invading species, while overlooking the substantial influence of the resident network’s structure on invasion success (Romanuk et al., 2009; Lurgi et al., 2014; Frost et al., 2019). The rationale behind this approach was that, given the primary objective of comparing the success of invaders in the presence and absence of each other, the nature of their ecological interactions and their respective positional indices could be the key factors. However, it’s essential to acknowledge that there exists a much broader range of species traits and network indices that can be explored (Romanuk et al., 2009; Móréh et al., 2021; Gouveia et al., 2021), and we plan to expand our research in this direction in future studies.

Thus, our study represents a first step towards a theoretical framework for understanding the relationships and consequences of both single and two-species invasions, a topic that has been extensively studied in the field and in mesocosms, but surprisingly little explored from a theoretical perspective.

## Supporting information

https://eltehu-my.sharepoint.com/:f:/g/personal/istvan_scheuring_ttk_elte_hu/Ep_vRDPyfTdOhVw1qhtU6YwBdBKSpkLMYixRzYPePWcqFg?e=fLBbkQ

## ACKNOWLEDGEMENTS

This work was supported by the European Union’s Horizon 2020 research and innovation programme under grant agreement No 952914. English grammar and style are checked by DeepL Write beta 2.0.

## Notes

### Competing Interest Statement

The authors have declared no competing interest.

