## Supplementary material for "Effects of joint invasion: how co-invaders affect each other’s success in model food webs?": https://eltehu-my.sharepoint.com/:f:/g/personal/istvan_scheuring_ttk_elte_hu/Ep_vRDPyfTdOhVw1qhtU6YwBdBKSpkLMYixRzYPePWcqFg?e=fLBbkQ

Ágnes Móréh, Ferenc Jordán, István Scheuring

**S1. Statistical analyses**

The association between the ecological relationships of the two invaders and the invasion outcomes was tested using the Chi square test. Due to the large sample size, it is not surprising that the results are significant. Since the number of categories in all cases was more than two, relevant differences that gave rise to significance were detected using the Bonferroni-corrected post-hoc z-test, with the Cramer's V effect size metric providing information on the strength of the relationships.

We applied the same approach to investigate whether the invasion outcome is influenced by the top or intermediate position of the invaders in the network. Additionally, we sought to determine which position is more likely to succeed in cases where prey and predator (all 1-step distance categories) or "lower" and "higher" trophic positions (trophic cascade and its combination with other categories) can be distinguished. We also employed a proportion (goodness of fit) test to examine whether the frequencies of the invasion outcomes, or the success of prey ("lower")/predator ("higher") species differed significantly from the overall probabilities observed in the whole sample.

The relationship between the topological network indices of the invaders was explored by analysing both the difference between the indices and their average. These two quantities are not independent of each other. A small difference between the indices can occur for both low and high averages, but as the difference between the two values increases, the average tends to move towards the mean. Thus, we did not attempt to express the relationship between the two values as a single measure; instead, we examined them separately. For differences, we used their absolute values. The extent to which the differences or means of the indices vary for different invasion outcomes were tested using a Welch ANOVA test because of the inequality of the variances of the samples. The variables were square root-transformed to ensure normality. Because of the significancy caused by the large sample sizes (Cohen, 1992), we took the effect size (𝜂2 and Cohen's d) into account to estimate the strength of the relationships. Since we compared multiple categories in both cases, we utilized a Games-Howell post hoc test to examine which group differences contributed to the significant results. To compare the indices of successful and unsuccessful invaders, we used Welch's t-test with Cohen's d effect size metric.

All of the statistical results are summarized in the Tables S2-S7. in the Supplementary Material.

**S2. Connection between the invaders' network indices or similarity and the invasion outputs (IOP)**

We capture the connection of the topological position of the two invaders by the difference (*d-'IND'*) and average of the network indices (*a-'IND'*) expressing them. We also investigate whether there is an effect of the invaders' similarity (*Jsim*) on the invasion outcome (the investigated outputs are detailed in Section 3.1 in the main text). We used 𝜂2 effect size metric for testing whether there is a significant difference between the differences and averages of these indices for each invasion output and, if so, how strong it is. The detailed results of the statistics can be found in the Supplementary Material (Table S4. and S5.). In general, it is true for all indices that their differences have a much smaller effect than their averages (𝜂2 obtained for *a-INDs* is always larger than for the *d-INDs,* see Table S4), i.e. the latter tend to determine the differences between the invasion outputs. However, this effect is small (𝜂2 < 0.06) or negligible (𝜂2 < 0.01) for almost all network indices. The number of the invaders' prey has the smallest effect on the invasion outcome (Fig. A1 e, 𝜂2 = 0.001), while their trophic level is the strongest influencing factor (Fig. A1 a, 𝜂2 = 0.09, see Table S4.).

The differences that contributed most to the overall - significant - effect were tested using a Games-Howell post-hoc test. Its results can be found in Table S5. and we have also highlighted the effect size values (Cohen's d) in Table S4, too. There are the largest differences between the IOP-categories in terms of the invaders' trophic level (*TL*). IOP-V (the case of complete failure) is the most different from the others (Fig. A1 a), with significantly higher a-*TL*s of the invaders, but also IOP-II (one invader fails) has higher a-TL with a medium effect size compared to IOP-I (Cohen's d = 0.64) and IOP-III (Cohen's d = 0.56). IOP-III and -IV are not considerably different, with small (|Cohen's d| < 0.5) or negligible (|Cohen's d| < 0.02) effect sizes. For all other network indices and the similarity of the two invaders, the difference between the invasion outputs is small or negligible and even not statistically significant, however, in case of betweenness centrality (*BC,* Fig. A1 b), IOP-V shows a moderate degree of divergence with respect to the others (except for IOP-II, where one invader also fails; Cohen's d = -0.29). Furthermore, for case IOP-V, in addition to a significantly higher a-*TL* and lower a-*BC* value, the invaders have a lower a-*CC* (Fig. A1 c, Cohen's d = -0.71), a-*D* (Fig. A1 d, Cohen's d = -0.66) and have less predators (a-*Npred*) (Fig. A1 f, Cohen's d = -0.61) than that of in the case of IOP-I.

However, according to Welch's test, the direction of the invaders' biomass changes investigated for IOP-I are not or only slightly influenced by their topological indices (see Fig. A2 and Table S4.). The overall effect size is negligible everywhere (𝜂^2^ < 0.01), the only small difference is revealed by the post-hoc test for betweenness centrality and similarity: slightly higher a-*BC (Fig. A2 b,* Cohen's d = 0.26*)* and *Jsim* (Fig. A2 h, Cohen's d = 0.29*)* are more likely to decrease the equilibrium biomass of both invaders.

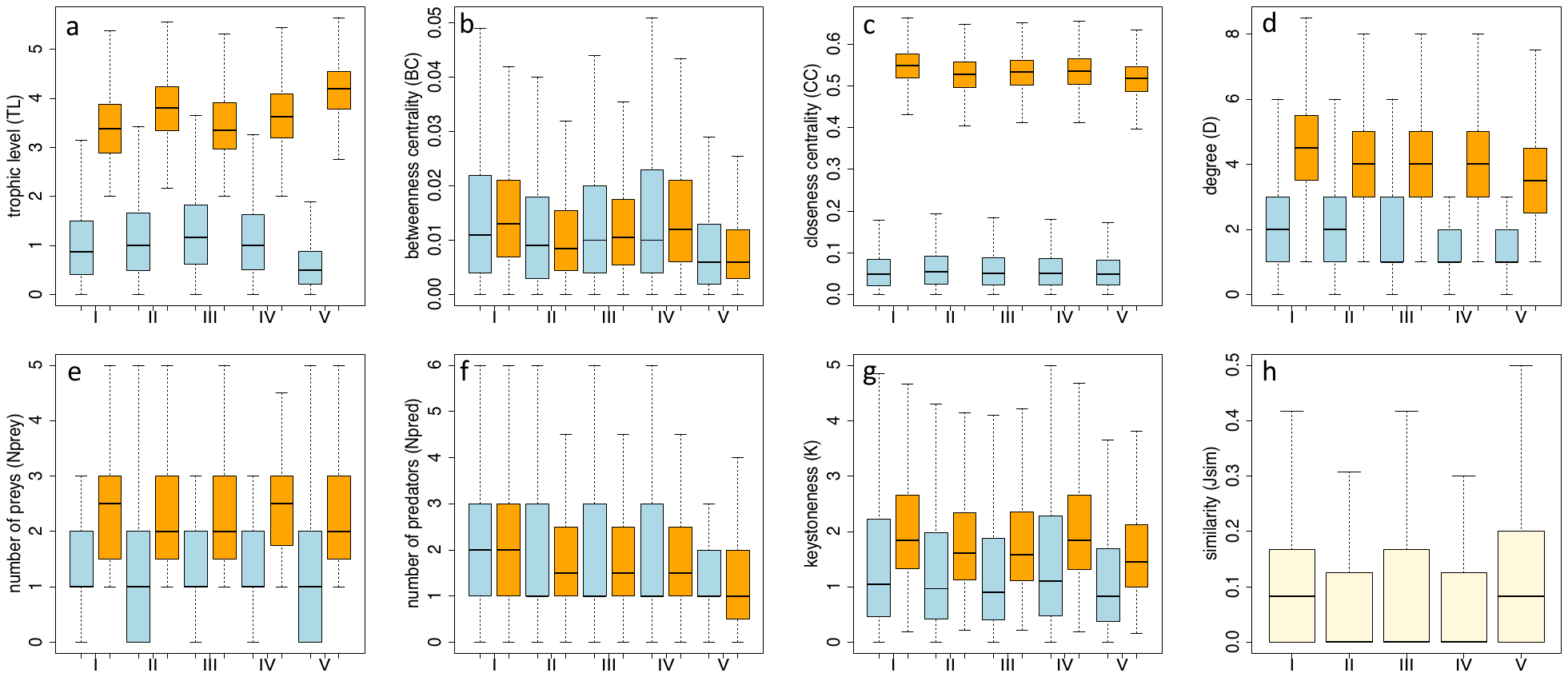

Fig. S1 The differences (blue) and the averages (orange) of the invaders' positional network indices (a-g) and similarity-values (beige, h), in case of the different invasion outputs (IOP), see Fig. 3. in the main text for the output categories.

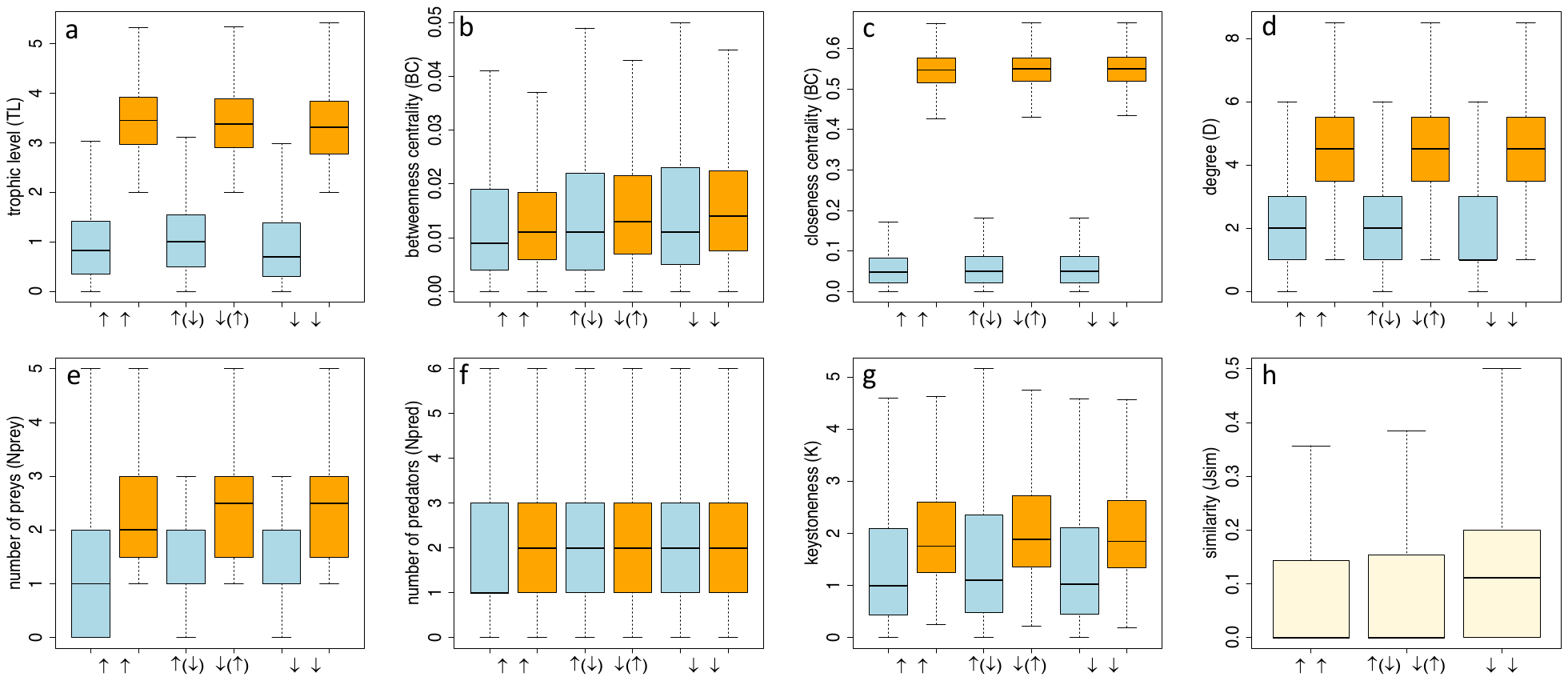

Fig. S2 Differences (blue) and averages (orange) of positional network indices (a-g) and similarity values (beige, h) of invaders for different directions of changes in their biomass in case of IOP-I (↑↑: both increase; ↑(↓) ↓(↑): one of them increases, the other decreases; ↓↓: both decrease).

**S3. Connection between the invaders' network indices and their success/failure**

We also investigated whether distinctions exist between the topological network indices of successful and unsuccessful invaders in cases IOP-II and IOP-III. The results of the Welch t-tests are presented in Table S7. The most informative metric is the effect size (Cohen's d) again, which effectively captures the magnitude of differences in indexes between successful (s) and unsuccessful (u) invaders. Our visual representations (Fig A3a-g.) clearly show that the most substantial difference is observed within the trophic level of the two invaders. Successful species consistently exhibit significantly lower TL values compared to unsuccessful invaders, irrespective of outcome type (IOP-II or -III, Cohen's d = -0.99 and -1.43, respectively). Conversely, higher BC-values are associated with a higher probability of invasive species' success, albeit with less significant impact (Cohen's d = 0.6 and 0.48 for IOP-II and -III, respectively). When considering closeness centrality, a significant but moderate distinction between successful and unsuccessful invaders is observed in case IOP-II (Cohen's d = 0.54), while for IOP-III, this distinction is negligible (Cohen's d = 0.17). A remarkably similar pattern emerges concerning the degree index (*D*). Notably, a moderate difference is evident for IOP-II (Cohen's d = 0.53), characterized by a higher probability of successful invaders having higher *D* values. Conversely, for IOP-III, the distinction in D indices is minor (Cohen's d = 0.22) according to invasion success or failure. While there is little difference in the amount of prey, successful invaders have a higher average number of predators in both outcomes (Cohen's d = 0.45 and 0.53 for IOP-II and -III, respectively). Keystoneness (K) reveals a small difference only in case of IOP-II, with no substantial distinction between successful and unsuccessful invaders in IOP-III. A higher K-value in case of IOP-II is more likely to be linked with a successful invader, but the effect size is small (Cohen's d = 0.29).

Interestingly, we do not see any difference in the topological network indices for biomass increase or decrease in case of IOP-I (Fig. A4a-g. and Table S7). Whether joint invasion increases its biomass or not, the trophic level of *I2* is always higher than that of *I1*. The same is true for the invaders’ number of predators, while the opposite is true for the number of preys: *I2* has significantly fewer preys than *I1*. It is true for all the indices investigated that whether the average of the groups analysed is slightly or significantly different, the differences are not correlated with an increase or decrease in biomass, but are consistently observed between the two invasive species.

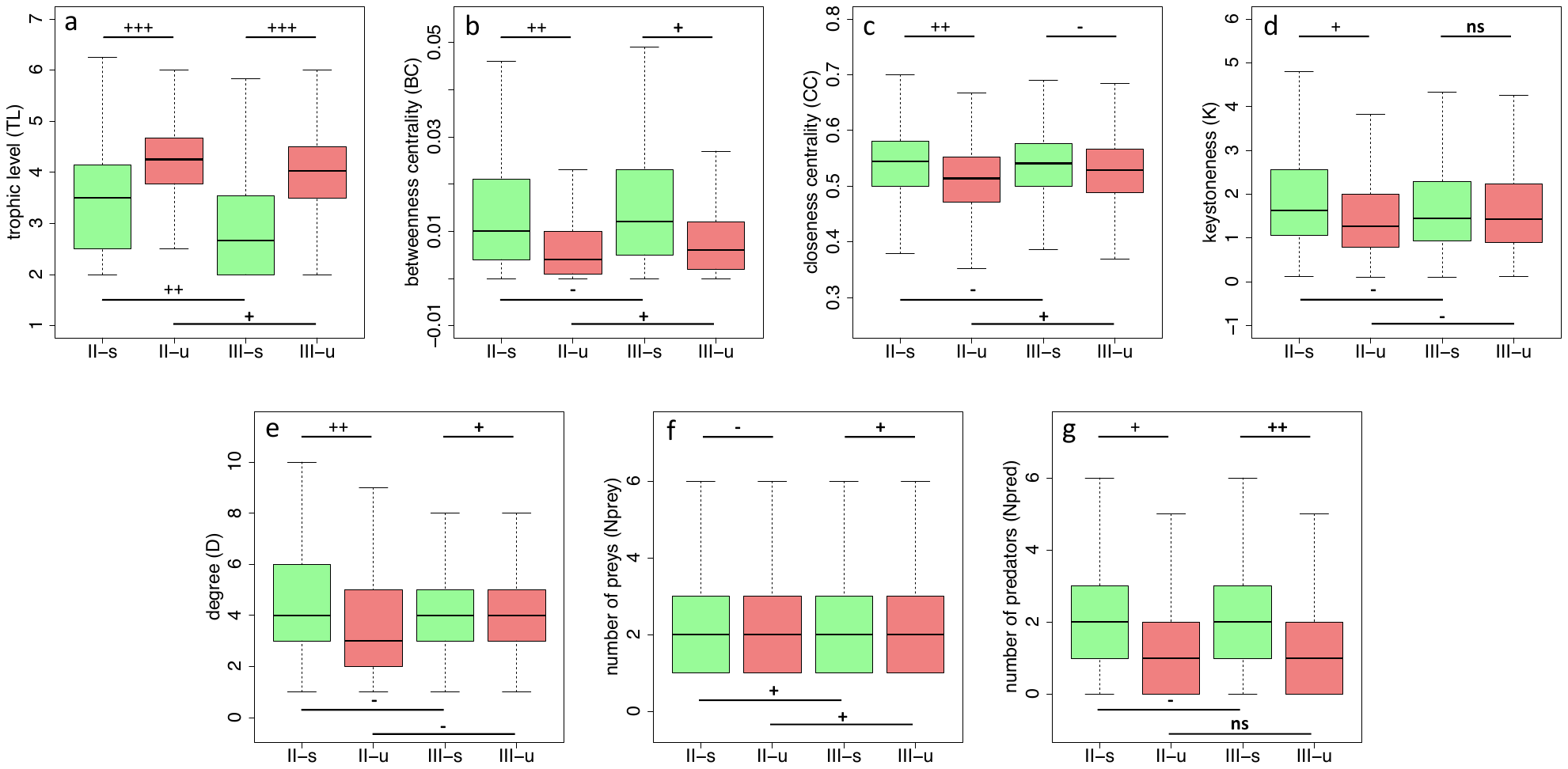

Fig. S3 Distributions of the successful (s, green) and unsuccessful (u, red) invaders' network indices in case of IOP-II and IOP-III. The lines above and below the boxplots represent the cases being compared. The strength of the relationship is indicated by Cohen's d effect size, which is categorized as non-significant (ns), negligible (-), small (+), moderate (++), or large (+++). For more details of statistics, see Table S7.

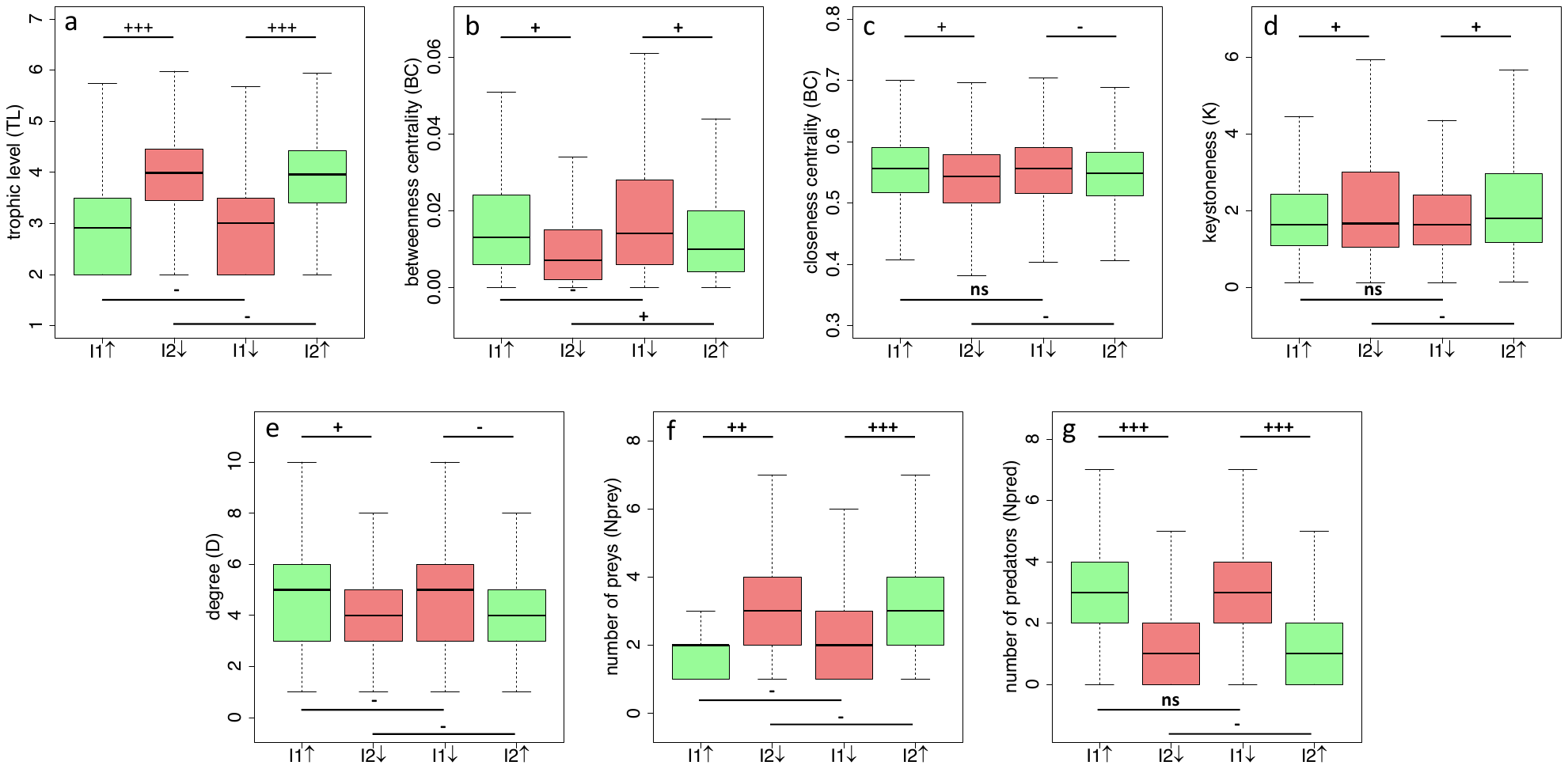

Fig. S4 Distributions of the topological indices specific to invaders with increasing (↑, green) and decreasing (↓, red) biomass in case of IOP-I. The lines above and below the boxplots represent the cases being compared. The strength of the relationship is indicated by Cohen's d effect size, which is categorized as non-significant (ns), negligible (-), small (+), moderate (++), or large (+++). For more details of statistics, see Table S7.
